# A comprehensive benchmark of discrepancies across microbial genome reference databases

**DOI:** 10.64898/2026.03.02.709103

**Authors:** Grigore Boldirev, Peace Aguma, Viorel Munteanu, David Koslicki, Mohammed Alser, Alex Zelikovsky, Serghei Mangul

**Author notes:** These authors jointly supervised this work (Mohammed Alser, Alex Zelikovsky, Serghei Mangul).

## Abstract

Metagenomic analysis of microbial communities relies significantly on the quality and completeness of reference genomes, which allow researchers to compare sequencing reads against reference genome collections to reveal essential community characteristics. However, the reliability of these analyses is often compromised by substantial discrepancies across existing reference resources, including differences in genome content, assembly fragmentation, taxonomic representation, and metadata completeness. While these inconsistencies are known to introduce bias, the extent of divergence between major databases remains largely unknown. Here, we present a comprehensive benchmark of discrepancies across multiple widely used microbial genome reference resources. We developed the Cross-DB Genomic Comparator (CDGC), which utilizes reference genome alignments to systematically capture discrepancies in genome assemblies across reference databases. Applying this framework, we found that 99% of viral genomes were identical across databases, indicating strong consistency in viral reference resources. In contrast, fungal genomes showed substantially greater variability: although 82% of assemblies exhibited at least 90% similarity, only 7% were identical across databases. More concerning, we identified a subset of 461 assemblies with less than 50% similarity, suggesting the presence of technical artifacts, incomplete assemblies, or damaged genome files that require closer examination. Collectively, these results demonstrate that systematic cross-database benchmarking provides a critical mechanism for refining the accuracy of individual reference databases and advancing efforts towards more unified and reliable universal reference genomes.

## Background

Metagenomics involves sequencing the DNA of samples that contain multiple organisms to explore the composition and functions of microbial communities^1,2^. This approach enables a comprehensive investigation of microbial composition, functional potential, and ecological interactions within various habitats. Advances in high-throughput sequencing technologies enable the generation of large volumes of reads, which, when matched against comprehensive reference genome databases, facilitate the detection and characterization of numerous microbial species and strains, including rare and previously unculturable organisms. Reference databases are crucial for accurate metagenomics analysis, providing the known genetic sequences, structural and functional annotations necessary to identify, classify, and understand the roles of microbes present in various habitats^3^. Reference databases are critical not only for taxonomic profiling^4^, but also for characterizing the metabolic pathways, gene functions, and ecological roles represented within microbial communities. By examining entire microbial communities, metagenomics may provide critical insights into microbial diversity, ecological functions, and the contribution of microorganisms to environmental processes and human health. Consequently, the reliability and completeness of reference databases directly influence the accuracy, reproducibility, and interpretability of metagenomic findings.

Current limitations of microbial reference databases pose substantial challenges to the accuracy and comparability of metagenomic research. These limitations include fragmented genome assemblies with high contig counts, incomplete or truncated sequence files, uneven representation of taxa, inconsistent or outdated taxonomic annotations, redundancy across databases, and the inclusion of low-quality genomes and metagenome-assembled genomes (MAGs) with variable completeness and contamination levels. While the impact of the choice of bioinformatics tool on metagenomics is well-documented, the influence of the underlying reference database is equally critical yet less standardized^5^. Researchers rely on these databases, which contain curated genome sequences and annotations, to accurately align sequencing reads, estimate taxon abundance, and assign gene functions. Yet, a significant portion of microbial diversity remains unrepresented, as even the most advanced tools cannot identify strains absent from the reference databases. Compounding this issue, previous studies have shown that database redundancy and taxonomic conflicts can severely degrade classification accuracy.^6^ Even well-characterized species can exhibit considerable variability across databases, differing in assembly quality, contig count, and taxonomic annotation.^7^ For instance, multiple genome assemblies may share the same taxonomic identifier despite major genomic differences—with one assembly being a complete genome and another a collection of fragmented contigs—or represent the same strain submitted over different years reflecting ongoing improvements in sequencing methods.^8^ These inconsistencies can lead to misaligned or misclassified reads, compromising taxonomic and functional profiles, hindering reproducibility, and introducing errors into comparative analyses.^5^ Consequently, the extent of error propagation from these database inconsistencies is likely underestimated, as few systematic evaluations have been performed.

To systematically identify and quantify divergences between major reference genome databases, we developed the Cross-DB Genomic Comparator (CDGC), a framework that uses reference genome alignments to capture inconsistencies among genome assemblies across reference databases and provides a precise, flexible, and reproducible way to quantify genomic similarity. Application of CDGC to bacterial (RefSeq^9^ and BV-BRC^10^), fungal (RefSeq and Ensembl^10,11^), and viral (RefSeq and Viral-Host DB^10–12^) reference collections revealed significant inconsistencies across databases. Nearly all viral genomes (99%) were identical across databases, and most fungal genomes (82%) shared at least 90% similarity. In contrast, bacterial genomes displayed a broader distribution of similarity values. Approximately half of bacterial genome pairs were completely identical across databases, while a comparable proportion showed similarity between 95% and 100%, and a smaller fraction exhibited similarity below 95%, indicating greater variability among bacterial assemblies overall. Notably, 461 assemblies exhibited less than 50% similarity across databases, consistent with potential technical artifacts or corrupted files. These discrepancies highlight the strong association between genome fragmentation and reduced similarity, emphasizing the importance of improving assembly completeness and database agreement. Ultimately, this work represents an important step toward improving the quality of individual databases and enabling the development of more comprehensive and standardized reference resources to support accurate and reproducible metagenomic analysis.

## Results

### Selected reference databases

Reference databases for this study were selected to represent widely used microbial genome resources with sufficiently structured metadata to support reliable comparisons across bacteria, fungi, and viruses. We extracted taxonomic identifiers (taxid), strain information, genome size, contig count, assembly release date, and FTP path for each entry, as these attributes enable both species-level identification and technical evaluation of genome assembly. To support robust cross-database comparisons of bacterial, fungal and viral genomes, we selected a curated subset of five primary databases—RefSeq^9^, BV-BRC^10^, Ensembl Fungi^11^, FungiDB^13^, and Virus-Host DB^12^—that provide broad taxonomic coverage and sufficiently structured metadata to ensure consistent and comparable analyses across domains. Several metadata attributes were essential for harmonization and assembly selection. The taxonomic identifier (taxid) provides a stable species-level reference that enables consistent matching across databases despite differences in naming conventions. Strain designation allows comparison of corresponding isolates when strain-level information is available. Genome size enables differentiation among multiple assemblies for the same strain and facilitates identification of the most complete assembly. Contig count allows stratification of similarity statistics and supports evaluation of the effect of assembly fragmentation on comparative results. Assembly release date enables selection of the most recent assembly when multiple versions exist. The FTP path provides a direct link to the corresponding FASTA file and ensures an exact connection between metadata and sequence data. Based on these criteria, we incorporated five reference resources into the study: RefSeq and BV-BRC for bacteria; RefSeq, Ensembl Fungi, and FungiDB for fungi; RefSeq andVirus-Host DB for viruses. Several additional databases were evaluated but excluded due to limitations in metadata completeness or structural consistency. JGI^14^ was excluded because strain information and, in some cases, taxonomic identifiers were inconsistently provided, preventing reliable harmonization, whereas UHGV^16^ was excluded because it consists exclusively of metagenome-assembled genomes, which are not directly comparable to isolate-based reference assemblies. GTDB^8^ was excluded due to difficulties in consistently linking downloaded genome FASTA files to their corresponding metadata entries, as some genome files lacked matching metadata records, and AllTheBacteria^17^ was excluded because key metadata fields, including strain designation, genome size, and assembly release date, were inconsistently reported. These criteria ensured that only databases with sufficient metadata completeness and traceability were included in the analysis. Metadata preprocessing and assembly selection were performed using metadata obtained on January 25, 2025. Metadata records were grouped by taxonomic identifier and strain designation to ensure that assemblies corresponding to the same strain were processed together. Within each group, assemblies were sorted by release date after conversion to a standardized datetime format. The most recent assembly was selected as the representative genome. When multiple assemblies shared the same release date, the assembly with the largest genome size was selected. This procedure ensured that a single representative assembly was chosen for each strain. After representative assemblies were selected, curated datasets were merged based on shared taxonomic identifiers and strain designations to identify corresponding genome pairs across databases. This process produced a harmonized dataset suitable for systematic comparative analysis. Genome assemblies were retrieved using accession identifiers and FTP paths provided in the metadata. For each genome, automated scripts constructed download URLs corresponding to the appropriate FTP location, including BV-BRC and NCBI servers. Genome FASTA files were downloaded, decompressed when necessary, and recorded in a log file to maintain traceability between metadata and sequence data. This automated retrieval framework ensured consistent and reproducible acquisition of genome assemblies for downstream comparative analysis.

### CDGC can systematically compare genome assemblies across multiple reference databases

The Cross-DB Genomic Comparator (CDGC) was developed to perform systematic, large-scale pairwise comparisons of genome assemblies across multiple reference databases. Our comparison strategy was determined by both metadata availability and the expected scale of biological variation within each microbial domain. For bacteria, strain-level identifiers were consistently available in database metadata, enabling direct strain-to-strain comparisons across resources. This resolution is biologically appropriate because genomic divergence among strains of the same bacterial species is typically modest relative to divergence observed between distinct species.^18^In contrast, viral and fungal databases lacked consistent and standardized strain-level annotations, precluding reliable strain-level matching across resources. For these groups, comparisons were therefore conducted at the species level. Although interspecies divergence generally exceeds intraspecies strain-level variation, the magnitude of genomic differences observed among viral and fungal species remains within a range that permits meaningful structural and sequence-level comparison, particularly when the objective is to assess cross-database assembly consistency rather than evolutionary distance. Using this approach, we compiled domain-specific datasets for systematic cross-database benchmarking, with strain-level resolution for bacteria and species-level resolution for viruses and fungi. Genomes with missing or unavailable sequence data were excluded. For multi-contig assemblies, contigs were concatenated in file order and treated identically to single-contig genomes. Genome similarity was determined by pairwise alignment of full genome sequences or concatenated contigs when genomes were fragmented. To ensure that equivalent assemblies were being compared across databases, we required matched genome pairs to have identical contig counts, thereby controlling for differences in assembly fragmentation. We summarized the distribution of shared bacterial genomes with matching contig counts from 1 to 10 in both databases (Supplementary Table S2). For bacterial genomes, we systematically evaluated contig-count concordance between RefSeq and BV-BRC and quantified cases in which the number of contigs differed between matched assemblies (Fig. S1). Genome pairs with mismatching contig counts were excluded from the similarity analysis so that comparisons were performed only between assemblies with the same number of contigs, ensuring that the same structural representation of each genome was being analyzed.

Whole genome alignment faces several significant challenges, particularly when dealing with large scale sequence data. Many alignment tools, such as AVID^19^, BLAST^20^, and FASTA^20,21^, were originally designed for smaller genomes, which makes them inefficient or unsuitable for large genomes or extensive datasets. Moreover, the correctness of sequence alignments produced by many existing tools has not been systematically evaluated, raising concerns about their accuracy, especially when applied to complex or highly variable genomes. To address this, we evaluated five whole genome alignment tools, including MUMmer4^20–22^, GSAlign^23^, DIALIGN-TX^23,24^, Progressive Cactus^23–25^, and BLAST, using a controlled artificial test case with known ground truth. The synthetic sequences were deliberately constructed to include predefined regions of perfect sequence identity, complete divergence, insertion and deletion events, and mismatched bases. Because the exact structure and positions of these regions were defined during sequence construction, the expected alignment outcome, including matches, mismatches, indels, and aligned and unaligned regions, was known in advance. The complete ground truth alignment and corresponding expected alignment statistics are provided in Supplementary Table S1.

Three tools, specifically BLAST, MUMmer4, and GSAlign, were successfully executed and evaluated, while DIALIGN-TX and Progressive Cactus were excluded due to specific technical limitations. DIALIGN-TX could not be executed because it required external configuration files and supporting parameter resources that were not available and could not be generated within the current computational environment. Progressive Cactus was excluded because it is designed for phylogeny-aware multiple genome alignment and requires a phylogenetic guide tree as mandatory input, which makes it unsuitable for direct pairwise alignment evaluation. A comprehensive summary of alignment statistics for all evaluated tools, including alignment lengths, matches, mismatches, indels, unaligned regions, and average identity values, is provided in Supplementary Table S1. BLAST precisely reproduced the ground truth alignment and achieved complete agreement with the expected alignment structure, as shown in Supplementary Table S1. MUMmer4 demonstrated high alignment accuracy and reported an average nucleotide identity of 97.99%, indicating strong concordance with the true sequence similarity. In contrast, GS Align showed substantially lower agreement with the ground truth and reported an average identity of 82.07%, reflecting overestimation of sequence variation and reduced alignment accuracy, as shown in Supplementary Table S1.

Based on these results, BLAST was selected as the primary alignment tool for all subsequent analyses. The alignment was executed using standard parameters, and the output was obtained in XML format to enable precise quantification of matches, mismatches, insertions, and deletions.

Although the tabular output format provides summary statistics for each alignment, it does not preserve positional information when alignments overlap, which can lead to double counting of mismatches and indels and inaccurate aggregate counts. To address this limitation, we used the XML output format, whose hierarchical structure allowed systematic parsing of all alignment segments and accurate computation of genome wide alignment statistics.

Many taxonomic IDs were not shared across databases Our comparison of major microbial genome databases reveals substantial discrepancies in the presence of reference genomes at the strain and species levels. These overlaps demonstrate that a researcher’s selection of a specific reference database significantly impacts the breadth of microbial taxa identified, potentially leading to varied results depending on the chosen resource. Analysis of bacterial records at the strain level comparing RefSeq and BV-BRC (PATRIC) reveals a total of 971,228 unique bacterial strains. While BV-BRC is more comprehensive, covering 94% (912,435) of the total strains identified across both platforms, RefSeq contains a distinct subset of 58,793 strains (6%) not found in the BV-BRC collection (Fig. 2a). When examining these bacterial resources at the species level, the combined set across Ensembl, RefSeq, and BV-BRC contains 117,288 unique bacterial species. BV-BRC represents the largest portion of this union with 84,108 species (approximately 72%), followed by RefSeq with 71,570 species (61%), and Ensembl with 20,261 species (17%). Notably, only 15,856 species are shared across all three bacterial databases, indicating significant divergence in species representation and suggesting that even large-scale aggregators may miss specialized or newly deposited data present in primary repositories.

**Figure 1.**
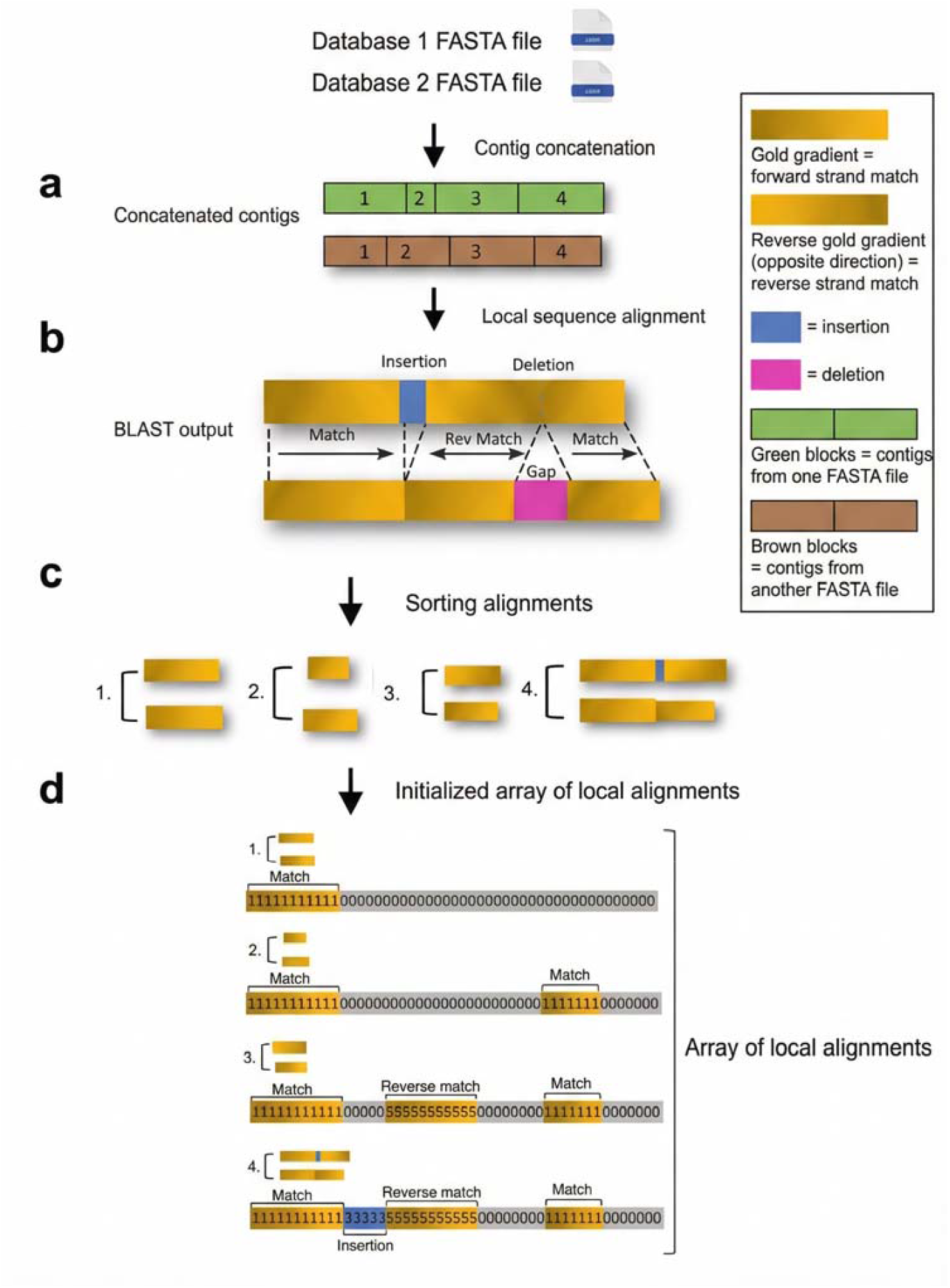
Cross-DB Genomic Comparator (CDGC) framework for array of local alignments. (a) Contigs from each genome assembly are concatenated in the FASTA file order to define a continuous coordinate system for both genomes. (b) Local sequence alignment is performed using BLAST to identify regions of similarity between query and subject genomes. (c) High-scoring segment pairs (HSPs) are extracted from the BLAST XML output and sorted by their positions in the subject genome. (d) Local alignment results are mapped position-by-position onto an array representing the subject genome. Each array position encodes the alignment outcome, including matches, mismatches, deletions relative to the query, insertions in the query, and strand orientation, while positions without alignment coverage remain unassigned. This positional encoding enables systematic quantification, storage, and downstream analysis of genome similarity at single-base resolution.

**Figure 2.**
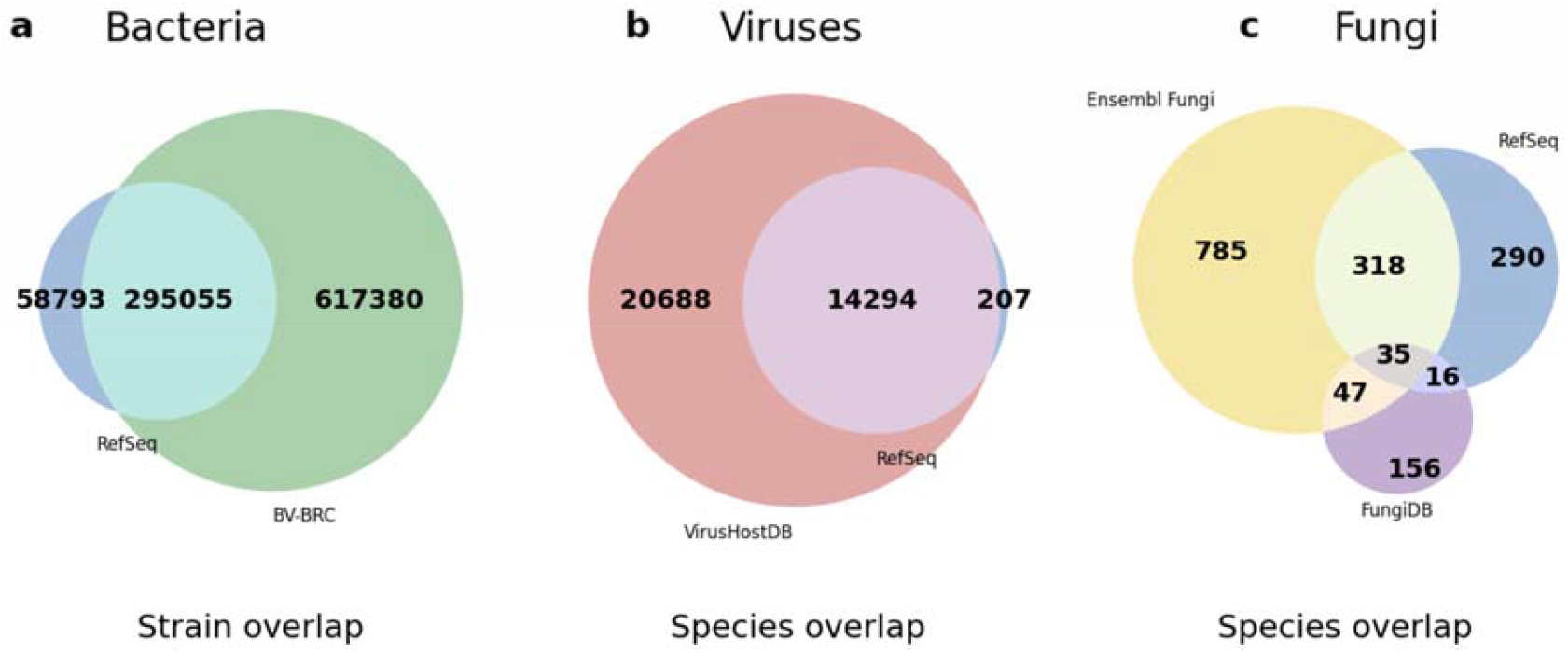
Overlap of taxa across major reference databases based on taxonomy IDs. Venn diagrams summarizing shared and unique taxa across bacterial, viral, and fungal reference resources. (a) Bacteria (strain level): Comparison between RefSeq and BV-BRC was performed at the species–strain level, where each element represents a unique species–strain pair with one representative assembly per database. Ensembl was not included due to incomplete and inconsistent strain-level annotations. (b) Viruses (species level): Comparison between VirusHostDB and RefSeq was performed at the species level, where each element represents a unique viral species present in the respective databases. (c) Fungi (species level): Comparison among Ensembl Fungi, RefSeq, and FungiDB was performed at the species level, where each element represents a unique fungal species represented in the respective databases.

A similar trend is observed in viral records at the species level. An analysis of RefSeq and VirusHostDB shows that across both resources, there are 35,189 unique viral species. VirusHostDB covers nearly 99% (34,982) of the total union, whereas RefSeq includes 14,501 species (41%). The two databases share 14,294 viral species, while VirusHostDB contributes a substantial number of unique entries that are absent from RefSeq (Fig. 2b). Despite the high level of coverage in VirusHostDB, RefSeq contributes a small but unique set of 207 species (0.6%) not indexed elsewhere. Differences are even more noticeable among fungal databases at the species level, where genomes are longer on average and strain-level information is often unavailable. Among the three fungal databases analyzed, we found 1,647 unique fungal species. Ensembl Fungi is the most comprehensive, accounting for 1,185 species (72% of the union), while RefSeq covers 659 species (40%) and FungiDB includes 254 species (15%). Only 35 fungal species are shared across all three databases, underscoring the limited overlap among fungal resources. Notably, RefSeq and FungiDB contribute 290 and 156 unique species, respectively, that are entirely absent from the other two databases (Fig. 2c).

The limited overlap between these repositories—particularly in the fungal kingdom—indicates that no single database currently provides an exhaustive catalog of microbial life. These findings suggest that relying on a single database for taxonomic assignment may result in the systematic omission of significant portions of known microbial diversity, which can skew downstream ecological or clinical interpretations. Consequently, researchers may need to consider multi-database integration or carefully evaluate the specific strengths of each repository relative to their target organisms to ensure robust and reproducible results.

### Complex contig alignment patterns across databases

As part of our cross-database genome similarity analysis, we randomly selected strains for manual examination to gain insight into matching patterns where species or strains are represented by multiple contigs. Upon comparing these assemblies, we found that the contigs did not align in a simple one-to-one manner. Instead, individual contigs from one assembly frequently exhibited complex alignments, either mapping to multiple contigs in the partner assembly or showing partial overlaps.

Comparison of the two assemblies for *Shewanella aestuarii* strain 17801 reveals two complex contig matching patterns across databases (Fig. 4a). RefSeq contig N9 aligns extensively with BV-BRC contig N9 across multiple discrete alignment blocks, with short internal regions of mismatches interrupting strongly matching sequence segments. At both ends of the aligned region, small segments are present that do not align to any counterpart, indicating short portions of sequence unique to each database. In addition, RefSeq contig N13 aligns to a leftward portion of BV-BRC contig N9 that is separate from the primary N9–N9 alignment. This indicates that sequence represented as two distinct contigs in the RefSeq assembly is incorporated within a single contig in the BV-BRC assembly. These observations demonstrate that differences in contig boundaries across databases generate complex matching patterns that are not apparent from contig numbering or from examining a single assembly alone. A similarly complex matching pattern is observed for RefSeq contig N2 and BV-BRC contig N2 (Fig. 4b). These contigs align across much of their shared length, although several localized mismatch regions interrupt otherwise strongly matching sequence segments. RefSeq contig N2 extends beyond the region aligning to BV-BRC contig N2, and its distal portion aligns with BV-BRC contig N5, forming a separate alignment block. Between the region aligning to BV-BRC contig N2 and the segment aligning to BV-BRC contig N5, RefSeq contig N2 contains an internal portion with no corresponding sequence in the BV-BRC assembly. This arrangement shows that sequence represented as a single contig in one database may correspond to multiple contigs in another and may also contain segments absent from the counterpart assembly. Together, these findings illustrate how differences in how genomic regions are divided into contigs across databases create complex cross-database matching patterns that become apparent through direct assembly-to-assembly alignment.

**Figure 3.**
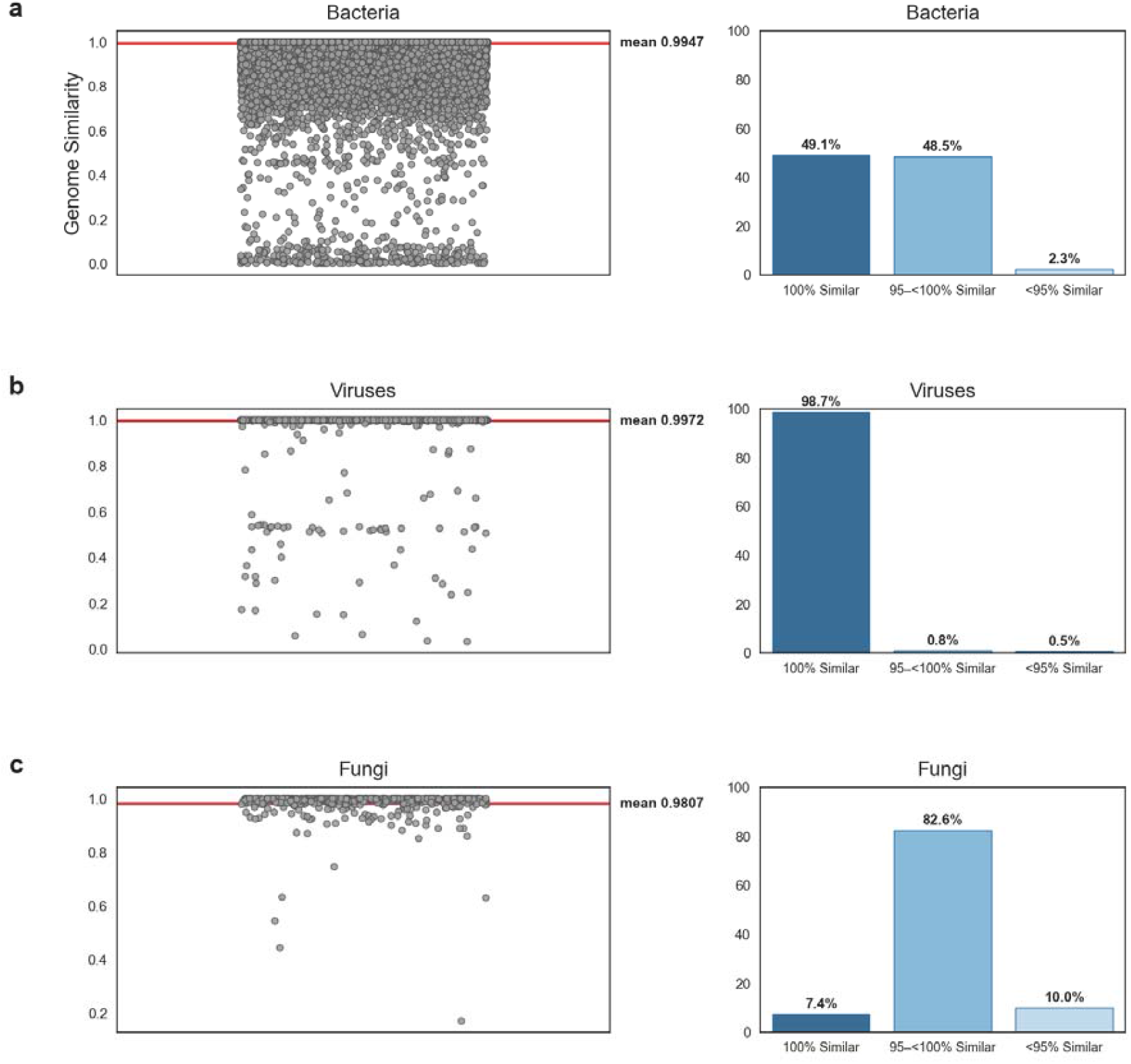
Genome similarity across reference databases for microbial genomes. For each microbial domain, the left panel shows the distribution of whole-genome similarity values for matched genome pairs (each dot represents one pair), and the right panel shows the proportion of pairs in three similarity categories (100%, 95–<100%, and <95%). (a) Bacteria, matched at the strain level. (b) Viruses, matched at the species level. (c) Fungi, matched at the species level.

**Figure 4.**
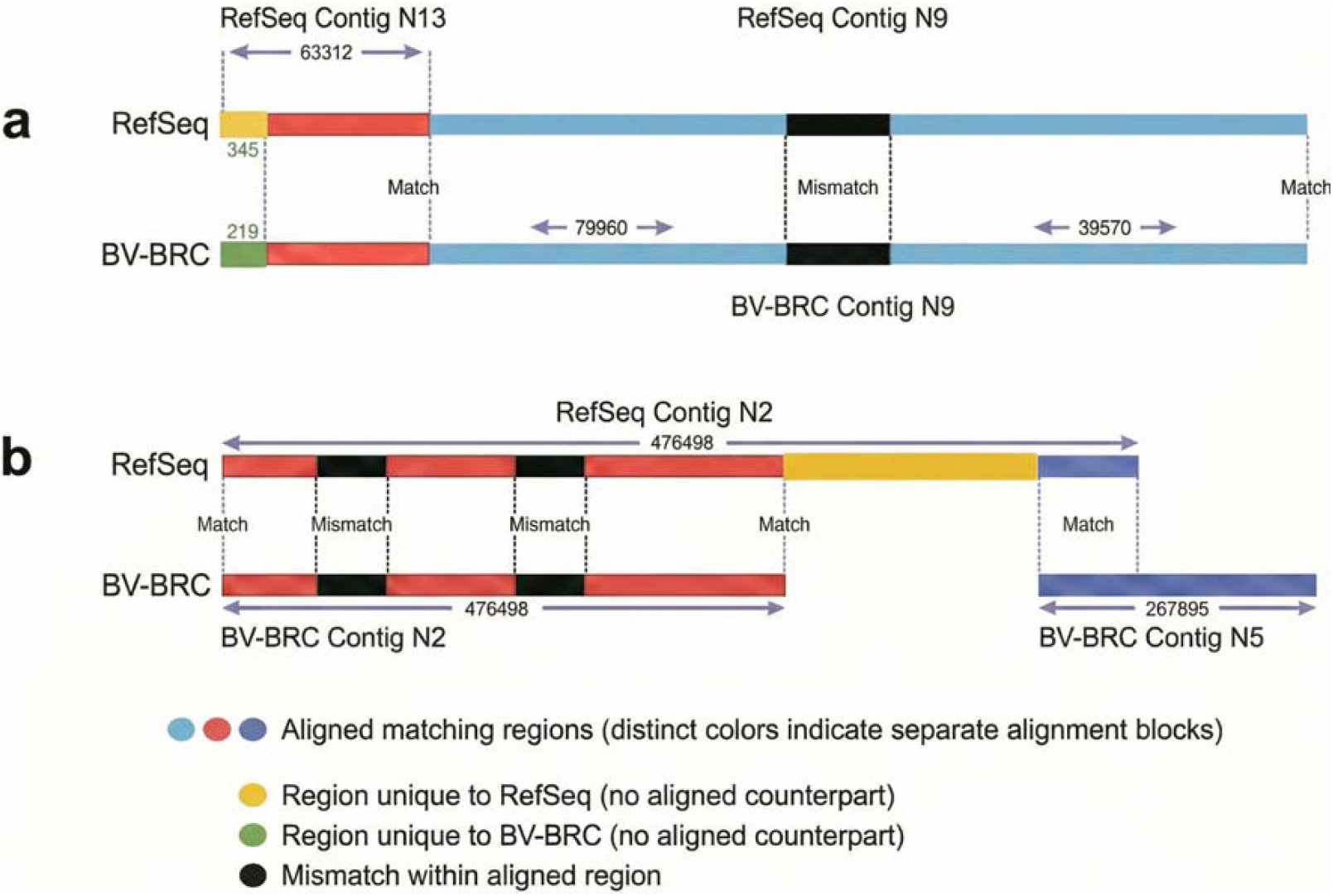
Alignment patterns between RefSeq and BV-BRC assemblies for *Shewanella aestuarii* strain 17801. (a) Comparison of contigs from two assemblies in RefSeq and BV-BRC. Both databases contain a contig labeled 9, and these contigs align across most of their length with localized mismatches. RefSeq contig 13 also aligns to BV-BRC contig 9 within the same genomic region shown for contig 9. (b) Comparison of contig 2 from RefSeq and contig 2 from BV-BRC. These contigs align across a substantial portion of their length with several mismatched segments. RefSeq contig 2 extends beyond the shared aligned region, and its terminal portion aligns with BV-BRC contig 5.

A recurrent pattern observed across species involves cases in which a single contig from one database aligns to two adjacent contigs from other database. This configuration is illustrated for *Acinetobacter guillouiae* strain KCTC 23200 (Fig. S3). The BV BRC assembly contains 100 contigs, whereas the RefSeq assembly contains 72 contigs, and the overall genome similarity between the two assemblies is low at 0.16. In this example, BV BRC contig 78 aligns almost entirely with two consecutive RefSeq contigs, 70 and 71. The left portion of BV BRC contig 78 aligns with RefSeq contig 70, and immediately downstream the alignment continues with RefSeq contig 71. The transition between these two alignments occurs with minimal unaligned sequence, indicating that RefSeq contigs 70 and 71 together represent a genomic region that is likely contiguous. This pattern shows that the sequence represented as a single contig in BV BRC is partitioned into two separate contigs in RefSeq. The two RefSeq contigs do not overlap with each other; instead, each aligns to adjacent segments of BV BRC contig 78. Their back to back alignment along contig 78 demonstrates that they cover neighboring regions of the same genomic locus. Such configurations reflect differences in contig fragmentation between databases and illustrate how assembly boundaries can divide what is represented elsewhere as a single contiguous sequence. Another interesting case observed within the same strain occurs when multiple contigs align to overlapping regions of the genome. For instance, contig 58 aligns partially with contig 36, while contig 1 also aligns to this same genomic region. The biological significance of these overlapping alignments remains uncertain; they could represent genomic duplications or other structural features. The length of the matching region shared among these three contigs is relatively small compared to the total genome length. (Fig S4) Other contigs from this species do not exhibit notable alignment patterns, and the overall genome similarity remains low. In these cases, individual contigs often show numerous short alignments with multiple contigs from the other assembly; however, these matches lack statistical significance due to their brevity. This pattern typically results in low overall alignment scores. For example, contigs 9 and 7 align with many different contigs, but the alignments are short and fragmented, contributing little to the overall similarity. Such fragmented and non-significant alignments are common among genome pairs with low similarity (Fig. S5).

For the strain *Leptospira interrogans* serovar str. LT1649, the assembly in BV-BRC comprises 874 contigs, whereas the corresponding RefSeq assembly contains 229 contigs. This discrepancy is commonly observed in genomes with low overall similarity: despite poor global alignments, short regions of similarity can occur that are represented in multiple contigs. The length of these regions is small relative to the total genome size. Biologically, such patterns may reflect duplicated genomic regions, repetitive sequences, assembly artifacts, or structural variations that hinder accurate contig resolution. (Fig.S6)

### Significant assembly discrepancies across microbial genome reference databases

Across microbial domains, the level of concordance between corresponding genome assemblies differed substantially, with viruses and fungi (matched at the species level) showing far higher consistency than bacteria (matched at the strain level). Viral genomes demonstrated the highest agreement across databases, with nearly all paired comparisons exhibiting complete or near complete similarity. Fungal genomes also showed strong concordance, with most assemblies aligning closely across sources. In contrast, bacterial genomes, which were matched specifically at the strain level to enable direct comparison, exhibited a much wider range of genome similarity values (Fig. 3).

Across microbial domains, viral genomes showed the highest level of agreement between databases (Fig. 3b). Among the 12,715 viral genome pairs analyzed, 98.7% showed 100% similarity, 0.8% showed 95% to less than 100% similarity, and 0.5% fell below 95%. The mean viral genome similarity was 0.9972. For fungi (Fig. 3c), 369 genome pairs were analyzed. Of these, 7.4% exhibited 100% similarity, 82.6% showed similarity between 95% and less than 100%, and 10.0% fell below 95%, with a mean similarity of 0.9807. When fungal comparisons were stratified by database source (Supplementary Fig. S7), the RefSeq–Ensembl pairs showed marginally stronger agreement (mean 0.9812) than the RefSeq–FungiDB pairs (mean 0.9790), although both distributions remained concentrated within the high similarity range. Bacterial genomes displayed substantially greater variability across databases (Fig. 3a). Of the 259,108 bacterial genome pairs evaluated, 49.1% exhibited 100% similarity, 48.5% showed similarity between 95% and less than 100%, and 2.3% fell below 95%, yielding a mean similarity of 0.9947. Although 97.6% of bacterial comparisons were at or above 95% similarity, the distribution exhibited a pronounced long tail of lower similarity values relative to viruses and fungi. It is important to note that the bacterial analysis included a much larger number of comparisons than the viral and fungal analyses (259,108 bacterial pairs versus 12,715 viral pairs and 369 fungal pairs), increasing the likelihood of detecting a broader range of similarity outcomes.

To assess whether genome similarity depends on assembly fragmentation, we focused on bacterial strains because they provided the most extensive contig count data. We performed balanced random sampling across contig count categories by selecting 5,187 genomes per category, corresponding to the smallest available group (assemblies with at least 500 contigs), and randomly sampling this number from all other categories. We then quantified the proportion of genome pairs falling into the 95% to less than 100% and below 95% similarity categories within each group. As shown in Supplementary Fig. S4, this normalized analysis did not reveal a consistent relationship between contig count and genome similarity, indicating that increased fragmentation alone does not systematically correspond to lower similarity estimates in this dataset.

### Cross-database comparison reveals incomplete genome assemblies

Our analysis revealed 461 strain pairs with genome similarity below 50%, indicating significant discrepancies at the assembly level between databases. Detailed examination of representative cases confirmed that these discrepancies were caused by incomplete, truncated, or low-quality genome sequence files rather than biological divergence.

One striking example involved the strain *Brachyspira hyodysenteriae* Bhyo_204, for which our framework reported approximately 50% genome similarity between the RefSeq and BV-BRC databases, suggesting a substantial discrepancy between the corresponding assemblies. To investigate this anomaly, we manually examined the BV-BRC genome sequence file and compared its length to the value reported in the metadata. The metadata specifies a total genome length of 3,122,276 bp across 8 contigs, whereas the downloaded sequence file contained only 1,524,611 bases. This means that 1,597,665 bases are missing, corresponding to more than half of the expected genome length. Thus, a substantial portion of the assembly sequence is absent from the file. This confirms that the reduced similarity observed by our framework was caused by an incomplete or truncated genome sequence rather than true biological differences. Such incomplete assemblies can severely distort similarity estimates and underscore the importance of validating sequence completeness when performing cross-database genome comparisons.

Another example involved the strain *Comamonas aquatica* NY8661, for which our framework reported an extremely low genome similarity ratio of 0.000395 between the RefSeq and BV-BRC databases, indicating a near-complete mismatch between the corresponding assemblies. Given that this genome is labeled as complete in the BV-BRC metadata, we manually examined the downloaded sequence file to determine the cause of this discrepancy. The metadata specifies a total genome length of 4,016,904 bp across 2 contigs, representing one chromosome and one plasmid. However, the downloaded file contained only 1,740 bases, meaning that nearly the entire genome sequence was missing. Further inspection confirmed that the file contained only the plasmid sequence, while the chromosomal sequence—the primary component of the genome—was entirely absent. Consequently, both the total sequence length and the expected number of contigs were inconsistent with the metadata. This confirms that the extremely low similarity ratio observed by our framework was caused by an incomplete genome sequence in BV-BRC, rather than true biological divergence, and highlights the importance of verifying both genome completeness and contig composition when performing cross-database genome comparisons.

A major discrepancy was also identified between the RefSeq and BV-BRC assemblies for *Bradyrhizobium* sp. UASWS1016 (taxon ID 1566379), despite both records originating from the same BioSample (SAMN03219997) and representing the same strain. The RefSeq assembly (GCF_001705105.1; ASM170510v1) has a total genome length of 7,960,052 bp, consistent with the expected genome size and gene content of *Bradyrhizobium* species. In contrast, the BV-BRC assembly (genome ID 1566379.4; GCF_000878315.1) is only 1,047,671 bp in length and consists of 107 contigs, representing only a small fraction of the expected genome content. Importantly, BV-BRC quality metrics explicitly classify this assembly as low quality, with a CheckM completeness score of 10.5% and contamination of 9.3%, and the genome quality flagged as “Poor” due to low completeness. These metrics indicate that most of the genome sequence is missing, confirming that the BV-BRC entry represents a severely incomplete and fragmented draft assembly rather than a complete genome. This large discrepancy in genome length and completeness explains the extremely low computed similarity between the two assemblies and demonstrates that the observed difference is caused by incomplete genome sequence data rather than true biological divergence.

## Discussion

We developed the Cross-DB Genomic Comparator (CDGC), a robust framework that encodes genome alignments to systematically capture base-level matches, mismatches, insertions, and deletions, enabling precise and reproducible quantification of genome similarity across reference databases. Application of CDGC revealed substantial inconsistencies among major microbial reference resources, highlighting a critical but underrecognized source of variability in genomic and metagenomic analyses. Viral genomes demonstrated near-complete concordance, with 99% of assemblies being identical between databases, likely reflecting their compact genome structure and lower structural complexity. Fungal genomes exhibited moderate variability, with 82% of assemblies showing at least 90% similarity, indicating generally high but not complete consistency across databases.

Accurate reference genomes are essential for high-resolution genomic analyses, and variation in genome representation across databases can influence sequence alignment, taxonomic assignment, and downstream interpretation. Even modest structural differences, such as fragmented assemblies or missing genomic regions, can alter similarity measurements and affect comparative analyses. Establishing a robust and scalable framework for systematic cross-database benchmarking is therefore critical to bridge these inconsistencies and guide future efforts toward harmonized reference resources. Integrating existing databases through structured comparative methodologies may enable the development of a more comprehensive and unified microbial genome reference, which is currently lacking but essential for reliable metagenomic analysis. Pangenome graph representations may offer a promising direction to organize this variation by placing multiple assemblies of a strain into one integrated graph, where shared segments align along the same path and differing segments are represented as alternative paths that branch away and later rejoin. This allows researchers to pinpoint where assemblies diverge, determine the extent of the divergence, and identify how and where they converge again. By examining all assemblies within a single graph structure, it becomes possible to clarify the origins of discrepancies between databases, identify regions that are absent or altered in specific resources, and understand how structural features vary across different assemblies of the same strain. Integrating these graph-based approaches can significantly enhance cross-database comparisons, improve interpretation of observed differences, and provide a stronger foundation for downstream metagenomic and comparative genomic studies.

Microbial reference databases, while foundational for open science, often prioritize aggregation over comprehensive curation.^7^. Our manual examination of low-similarity assembly pairs, such as the case of Shewanella aestuarii strain 17801, highlights the unique opportunity to merge the contigs from individuals databases into longer contigs or potentially complete genome. Specifically, BLAST alignment analysis revealed that RefSeq contig 9 aligns with PATRIC contig 9, indicating that these contigs represent the same genomic region (Fig. 4). However, RefSeq contig 13 also aligns to the same PATRIC contig 9 and overlaps with the corresponding region of RefSeq contig 9, demonstrating that contigs 13 and 9 in RefSeq represent adjacent or overlapping segments of a continuous genomic region that could have been merged into a single contig.

To further investigate extreme discrepancies, we manually analyzed all cross-database assembly pairs exhibiting less than 50% genome similarity, which resulted in a subset of 461 cases for detailed inspection. The 461 cases identified by CDGC collectively demonstrate that extremely low genome similarity values are primarily associated with missing or incomplete genome sequence files, truncated assemblies, or low completeness draft genomes. During manual inspection of these cases, we also observed that a subset of assemblies had already been annotated by the source databases as low quality or incomplete. These quality designations were not incorporated into the initial comparative filtering criteria, as our analysis focused on cross database sequence concordance independent of database specific quality labels. Consequently, some of the extreme discrepancies reflect assemblies that were previously flagged as suboptimal by their respective repositories. This represents a limitation of the current analysis and underscores the importance of integrating standardized assembly quality metadata into large scale cross database comparisons.

Our findings confirm that cross database similarity analysis can serve as an effective method for identifying assembly completeness issues and highlight the importance of validation when performing large scale comparative genomic analyses across reference databases. Establishing consistent standards for reference database development remains essential for the design and validation of bioinformatics tools and pipelines. Limited curation can reduce reliability and impair comparability across studies. Addressing these challenges will require coordinated efforts among database providers to reconcile discrepancies and develop unified strategies for data integration and automated curation. By systematically applying tools such as CDGC and incorporating complementary approaches including pangenome graph methods, potential errors can be more effectively detected and resolved, thereby improving overall genome quality.

## Methods

### Database Selection

We aim to use microbial databases that are both widely recognized within the research community and rich with sufficiently structured metadata to enable reliable comparisons across bacteria, fungi, and viruses. We prioritized metadata fields that are relevant for this study, such as the taxonomic identifier (taxid), strain name, genome size, contig count, assembly release date, and FTP path for downloading the genome sequence. These fields together provide the information needed to determine the species represented, identify the strain when such information is available, and evaluate the characteristics of each assembly.

We utilized a total of six reference databases across the bacterial, fungal, and viral domains. (Table 1.) The databases were: two bacteria databases, RefSeq and BV-BRC, three fungi databases, RefSeq, Ensembl Fungi, and FungiDB, and two virus databases, RefSeq and Virus-Host DB. Several other databases were reviewed during the initial assessment but were not incorporated into the analysis due to characteristics that made them less suitable for the goals of this study. For example, we observe that the JGI database does not consistently provide strain-level information and, in some cases, lacks complete species-level assignments. We observe that the UHGV database contains only metagenome-assembled genomes (MAGs), which represent environmental or metagenomic sequences rather than isolate-based reference genomes. We also observe that the GTDB database requires additional processing steps to reliably associate its downloaded genome FASTA files to their corresponding metadata. In our preliminary analysis, this association was not consistently successful: some genomes could not be matched to any metadata record, either because the relevant files were absent from the downloaded dataset or because inconsistencies were introduced during our own processing pipeline. We also observe that the AllTheBacteria database does not consistently provide fields such as strain designation, genome size, or assembly date, and some of its tables contain structural inconsistencies that limit reliable processing. These challenges introduce uncertainty when processing large databases, which highlights the dire need for more standardized and robust association between genome files and their metadata.

**Table 1.** Overview of major genomic databases used in this study, including RefSeq, BV-BRC, Ensembl, the Virus-Host Database, and RefSeq subsets for viruses and fungi. The table summarizes each database’s description, key bacterial, viral, or fungal statistics derived from relevant metadata fields (e.g., assembly_level, genome_status, taxid), and provides direct links for data access.

| Database | Primary Focus / Data Type | Scope (Species/O rganisms) | Key Strengths / Unique Features | Access Methods | Number of species | Number of strains |
| --- | --- | --- | --- | --- | --- | --- |
| **RefSeq** | Curated genomes, transcripts , proteins | Bacteria, Viruses, Fungi | High-quality, standardized, NCBI-managed, foundational resource. | Website , FTP, BLAST | 83490 | 329565 |
| **BV-BRC (PATRIC)** | Bacterial genomic, proteomic, clinical, environmental data | Bacteria | Comprehensive bacterial data, integrates PATRIC, functional annotation. | Website , FTP, API | 104336 | 878074 |
| **ENSEMBL** | Genomic data (genes, features, sequences ) | Bacteria, Archaea, Fungi | Flexible access (exports, REST API, MySQL dumps), detailed genomic. | Website , REST API, FTP | 20325 Bacteria 1185 Fungi | Not applicable |
| **Virus Host DB** | Virus-host interactions, viral genomes, host info | Viruses & their hosts | Specialized interaction data, manually curated literature. | Website | 34982 | Not applicable |
| **FungiDB** | Fungal/oomycete genomic & functional data | Fungi, Oomycetes | Interactive platform, metabolic pathways, gene functions, part of VEuPathDB. | Website | 254 | 331 |

### Standardizing the taxonomy across the microbial reference databases

Processing strains in microbial reference databases is challenging because existing strain identifiers cannot serve as reliable unique keys for linking reference genomes to their corresponding metadata. Strain identifiers lack standardized numerical labels and are generally recorded as free-text fields, leading to substantial variability across data sources. Ideally, we need a unique strain identifier that provides a stable strain-level identifier and helps minimize ambiguity even when strain names are formatted differently across databases. To match records across the microbial reference databases, we generate strain-level identifiers using strain information that are usually stored in metadata. Each database provides this information in a different format, so the appropriate column in the metadata has to be carefully identified for each database. For example, in RefSeq the strain designation is listed in the infraspecific_name column using the format “strain=…”, from which we extract the substring following the equal sign. Other databases include a dedicated strain column or a comparable attribute containing the submitted strain label, which we processed in an analogous manner. After extracting these text-based strain names, we performed exact string matching across the metadata tables to identify corresponding strains shared between databases. This workaround approach allows us to detect strain-level overlap but is inherently limited because strain names are not standardized identifiers and are entered manually by contributors. As a result, inconsistencies in spelling, formatting, or naming conventions can affect matching accuracy. Despite this limitation, text-string comparison remains the only feasible strategy for harmonizing strain information across large, heterogeneous microbial reference databases.

### Cross-DB Genomic Comparator (CDGC)

Different reference databases often use the same strain name, yet they may contain slightly or even substantially different versions of the corresponding reference genome sequences. This necessitates the need to develop a genome-matching approach that can determine whether sequences are identical and, if not, quantify the extent of their differences. We evaluated state-of-the-art whole genome alignment tools such as MUMmer4^22^, GSAlign^23^, DIALIGN-TX^24^, Progressive Cactus^25^, and BLAST. BLAST was the only tool that precisely provided an alignment matching the ground truth alignment (Supplementary Table S1). However, BLAST provides local alignments between subsequences of genomes, where an alignment of two subsequences may overlap with the alignment of another two subsequences without providing positional information. This can lead to double counting of mismatches and indels and inaccurately aggregate counts.

We developed a novel, high-accuracy approach for base-level genome comparison. We exploit the output of BLAST to build a consensus of all subsequence alignments by capturing both the coverage and the structural differences across the entire genome. This allows consistent quantification of matches, mismatches, and insertion–deletion events while preserving their exact locations in the subject genome. Our CDGC framework starts by examining the two input FASTA files, one from the first database (query) and another from the second database (subject). If a FASTA file has several contigs, we concatenate all of them into a single contig to ensure that each genome is represented in a single coordinate system, allowing direct positional mapping of alignment results (Fig. 1a). We then apply BLAST to the two input FASTA files to produce a local alignment (in an XML format) between many two subsequences of the concatenated genomes (Fig. 1b). The BLAST XML output is parsed to extract all alignments, called high-scoring segment pairs (HSPs), defining regions of similarity and difference between the two subsequences. These extracted alignments are sorted according to their percentage identity and length. The percentage identity measure is calculated by dividing the number of exact matches in an alignment by the length of the alignment; gaps are counted as mismatches. This ordered structure is critical to ensure that the framework prioritizes long alignment segments over random short alignments, and consistently labeling the alignments when building the consensus of all local alignments (Fig. 1c). To generate the alignment consensus, we initialize an array whose length equals the length of the subject genome sequence (Fig. 1d). Each position in the array stores an integer (ranging from 0 to 7) encoding the alignment outcome at that location. Positions not covered by any HSP pairs remain 0, indicating no alignment coverage. A value of 1 represents a match by direct HSP, while 5 represents a match by a reversed HSP (i.e., an alignment of its reverse complemented sequence). A value of 3 denotes a position deleted from the query sequence, while 7 indicates a deletion happened in its reverse complemented sequence. A value of 4 indicates a mismatch between subject and query in this corresponding position. When there is a gap in the subject (a base exists in the query, but missing from the subject), the value at the position after which gap occurs is negated (for example, 1 becomes −1, or 5 becomes −5), indicating the gap occurs between current and next position. This approach preserves positional accuracy and structural context while keeping previously established labels.

After populating the array, we count the occurrences of matches, mismatches, insertions, and deletions and record these summary statistics along with metadata in an output CSV file. The array is then serialized to a binary file for efficient storage and downstream analysis. This positional encoding enables comprehensive and reproducible genome comparison while preserving the full structure of alignment relationships between query and subject genomes. To effectively capture both nucleotide level agreement and completeness of alignment across the genome, and provide a global assessment of assembly concordance, we develop a genome similarity measure, defined as the number of matches and reverse matches divided by the full length of the subject genome. This measure is derived directly from the alignment array produced by the CDGC framework, which encodes alignment outcomes for every base position in the subject genome after concatenating contigs into a single continuous sequence. Because the denominator is the full genome length, this metric reflects agreement across the entire assembly, including both aligned and unaligned regions. Positions that do not align due to missing sequence, structural differences, or assembly inconsistencies remain explicitly represented and reduce the similarity score.

A commonly used alternative measure of genome similarity is Average Nucleotide Identity (ANI), which quantifies the average nucleotide identity between homologous regions shared by two genomes.^26^,^27^ ANI is typically computed by identifying aligned fragments between genomes, calculating nucleotide identity within each aligned segment, and averaging these values to produce a genome level estimate. Because ANI is calculated over aligned regions, portions of the genome that do not align may be excluded or contribute less to the final value. Consequently, ANI reflects sequence similarity within homologous regions, whereas the CDGC similarity measure reflects agreement across the entire genome by incorporating both identity and alignment completeness into a single value.

### Metrics

From each CDGC alignment array, we extracted the counts of matches, mismatches, insertions, and deletions, which collectively provide a complete positional characterization of the alignment between two assemblies. These values, together with the alignment arrays themselves, were systematically recorded and serialized to ensure reproducibility and to retain a permanent quantitative record for every genome pair. Because these quantities fully define the alignment structure, the framework permits computation of genome similarity metrics directly from the alignment representation. In this study, genome similarity was defined as

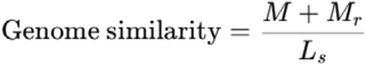

where MM is the number of matches in forward orientation, MrMr is the number of matches in reverse orientation, and LsLs is the total length of the subject genome. This definition ensures that similarity is normalized by the full genome length and can be computed directly and reproducibly from the alignment array. Although other similarity measures can be derived from the same representation, this metric provides a consistent and straightforward measure of genome level agreement across assemblies.

### Comparing assemblies with multiple contigs

Some genomes can be labeled as “complete” in the metadata but are represented by two or more contigs. To analyze these genomes consistently, we concatenated the contigs in the order they appeared in the original assembly file, effectively creating one longer sequence per genome. This allowed us to apply the same analysis pipeline to both multi-contig and single-contig genomes without modification. We then experimentally compared the results of aligning a query genome to either concatenated contigs or to a single-contig genome and found that BLAST alignments were consistent between the two approaches. This consistency confirms that concatenating contigs does not introduce bias or distortions in the similarity comparisons, enabling reliable analysis across different assembly types.

### Visualization

For the alignment cases highlighted in the Supplementary Material to illustrate matching patterns across assemblies, visualizations were generated using ALVIS, an interactive non-aggregative tool for exploratory analysis of multiple sequence alignments.^28^

---

**Figure S1.**
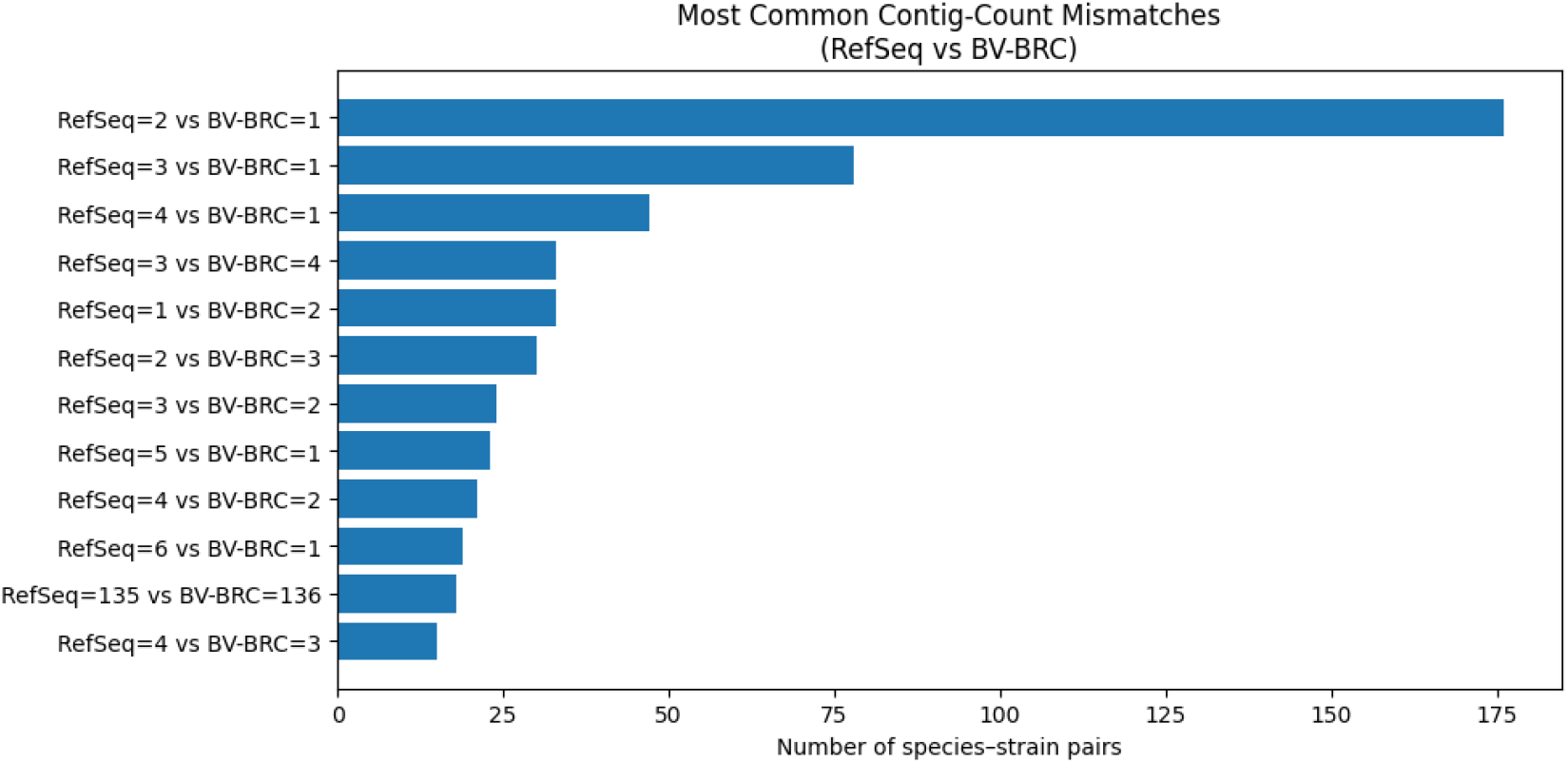
Most common contig-count mismatches among shared species–strain pairs between RefSeq and BV-BRC. The bar chart summarizes the most frequent cases in which representative assemblies matched at the species–strain level exhibit different contig counts across the two databases, with a total of 10,752 such mismatched pairs. Each bar corresponds to a specific RefSeq– BV-BRC contig-count combination, ordered by frequency. These assemblies were excluded from the genome similarity analysis to ensure consistent and comparable alignment conditions between matched genome pairs.

**Figure S2.**
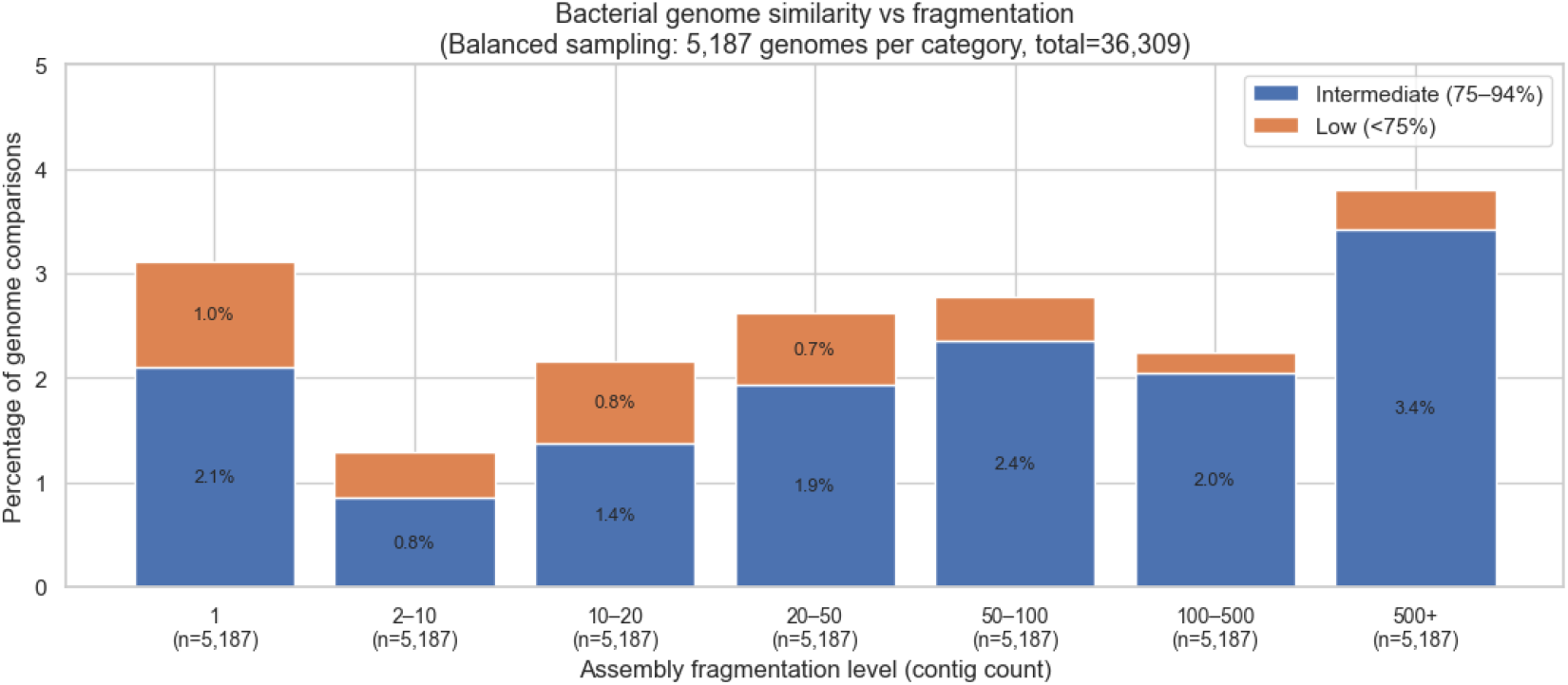
Distribution of genome similarity scores across bacterial strains from the BV-BRC and RefSeq databases with varying levels of contiguity. Stacked bars show the proportion of intermediate, and low similarity comparisons for genomes grouped by assembly fragmentation level.

**Figure S3.**
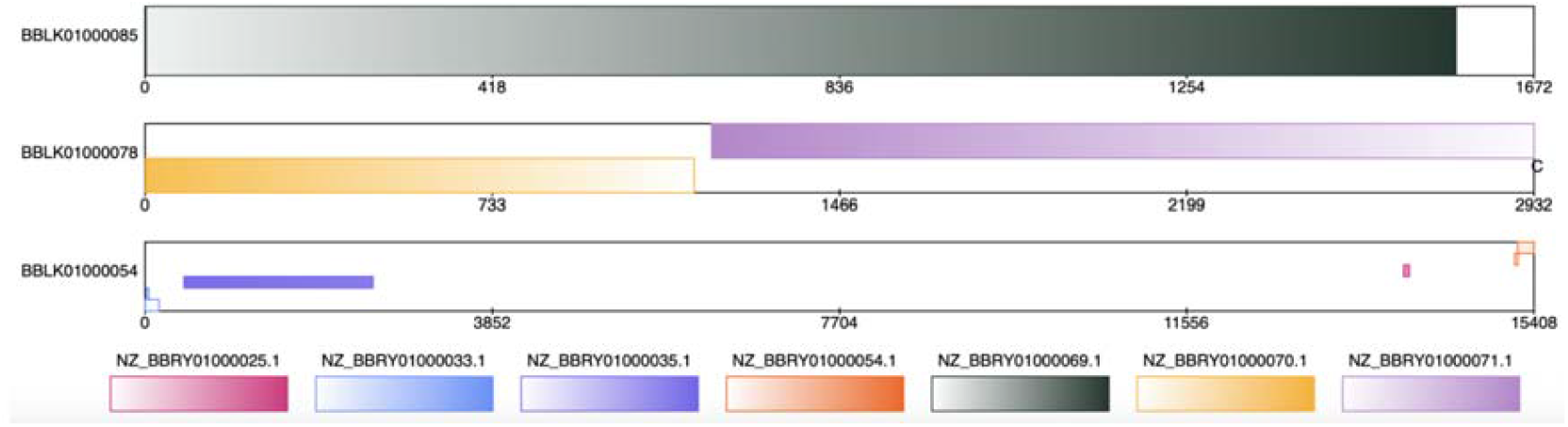
Adjacent contig partitioning in Acinetobacter guillouiae strain KCTC 23200. Assembly to assembly alignment shows that BV BRC contig 78 corresponds sequentially to two consecutive RefSeq contigs, 70 and 71. The two RefSeq contigs align back to back along contig 78 with minimal interruption, indicating that a continuous genomic region in BV BRC is represented as two separate contigs in RefSeq. This configuration illustrates database specific differences in contig fragmentation despite low overall genome similarity.

**Figure S4.**
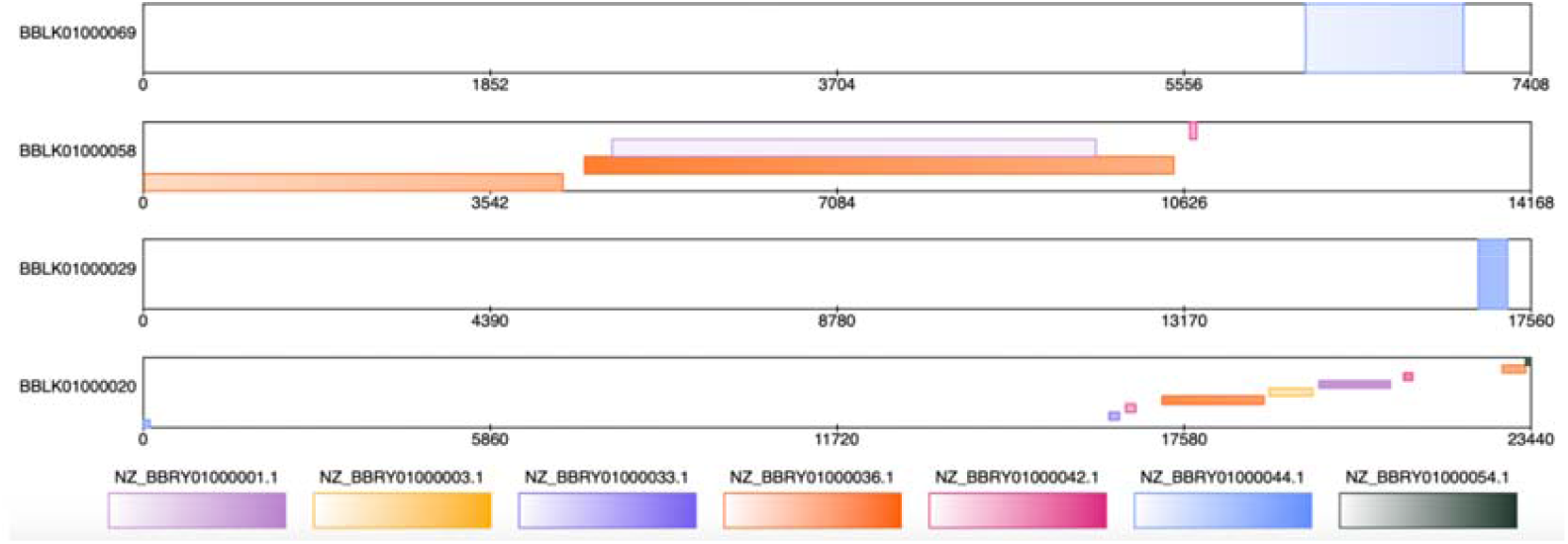
Overlapping contig alignments within Acinetobacter guillouiae strain KCTC 23200. Multiple contigs map to partially overlapping regions of the same genomic locus. Contig 58 and contig 36 share an overlapping aligned segment, and contig 1 also aligns within this region. The shared aligned portion is short relative to the genome length, suggesting localized duplication, repetitive sequence, or assembly related effects rather than large scale concordance.

**Figure S5.**
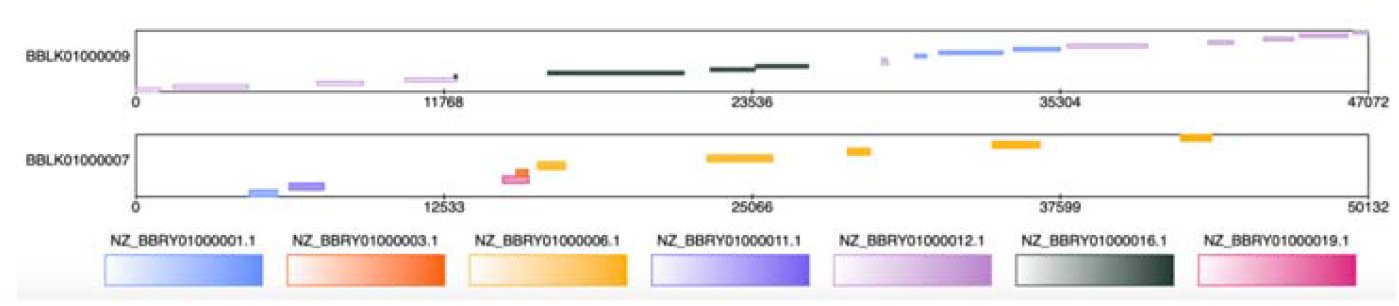
Fragmented short alignments associated with low genome similarity. Representative example of assemblies in which numerous brief alignment blocks occur between many contigs, but without extended contiguous matches. Although contigs such as 9 and 7 align to multiple counterparts, the alignments are short and dispersed, contributing minimally to total similarity and resulting in low overall alignment scores. Comparison of a highly fragmented BV BRC assembly (874 contigs) with a less fragmented RefSeq assembly (229 contigs) reveals sparse, short alignment segments scattered across contigs. Aligned regions are small relative to genome size and occur in multiple locations, consistent with limited global similarity and complex structural correspondence between assemblies.

**Figure S6.**
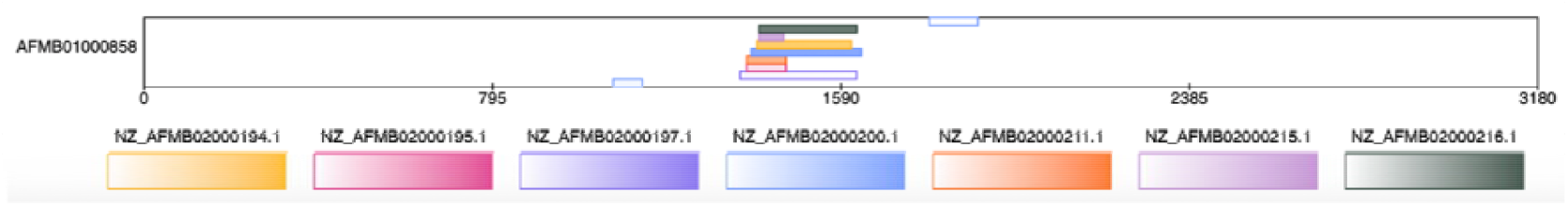
Extensive contig fragmentation in *Leptospira interrogans* serovar LT1649. Comparison of a highly fragmented BV BRC assembly (874 contigs) with a less fragmented RefSeq assembly (229 contigs) reveals sparse, short alignment segments scattered across contigs. Aligned regions are small relative to genome size and occur in multiple locations, consistent with limited global similarity and complex structural correspondence between assemblies.

**Figure S7.**
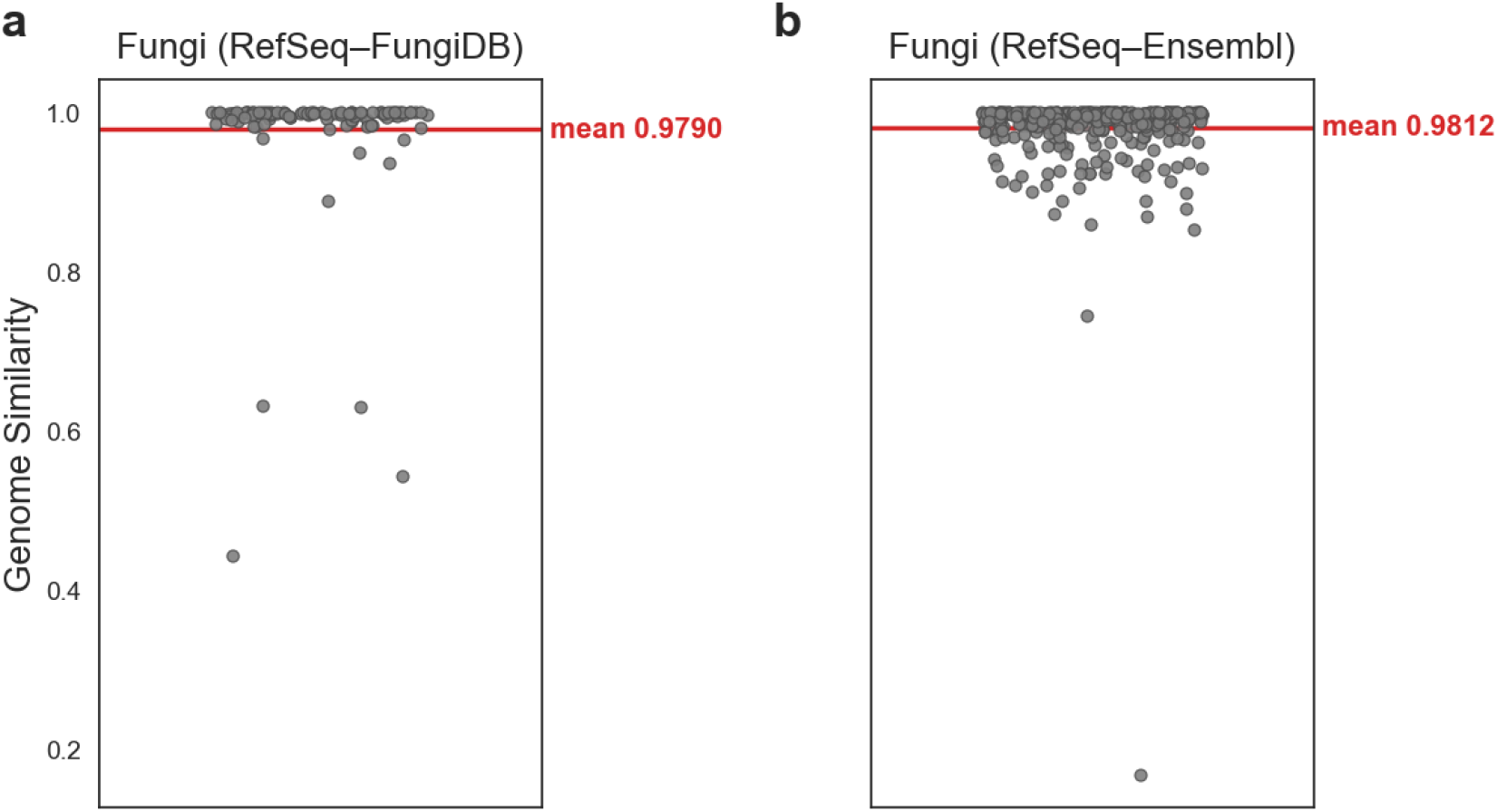
Genome similarity among fungal species. Each point represents a single pairwise comparison between assemblies of the same fungal species obtained from two independent reference databases. Panel a shows comparisons between RefSeq and FungiDB, and panel b shows comparisons between RefSeq and Ensembl. The y axis indicates genome similarity calculated for each species level comparison. The red horizontal line marks the mean similarity across all comparisons within each dataset. Most species exhibit near complete agreement between databases, although a subset of comparisons shows substantially lower similarity, indicating discrepancies in genome representation across resources.

**Figure S8.**
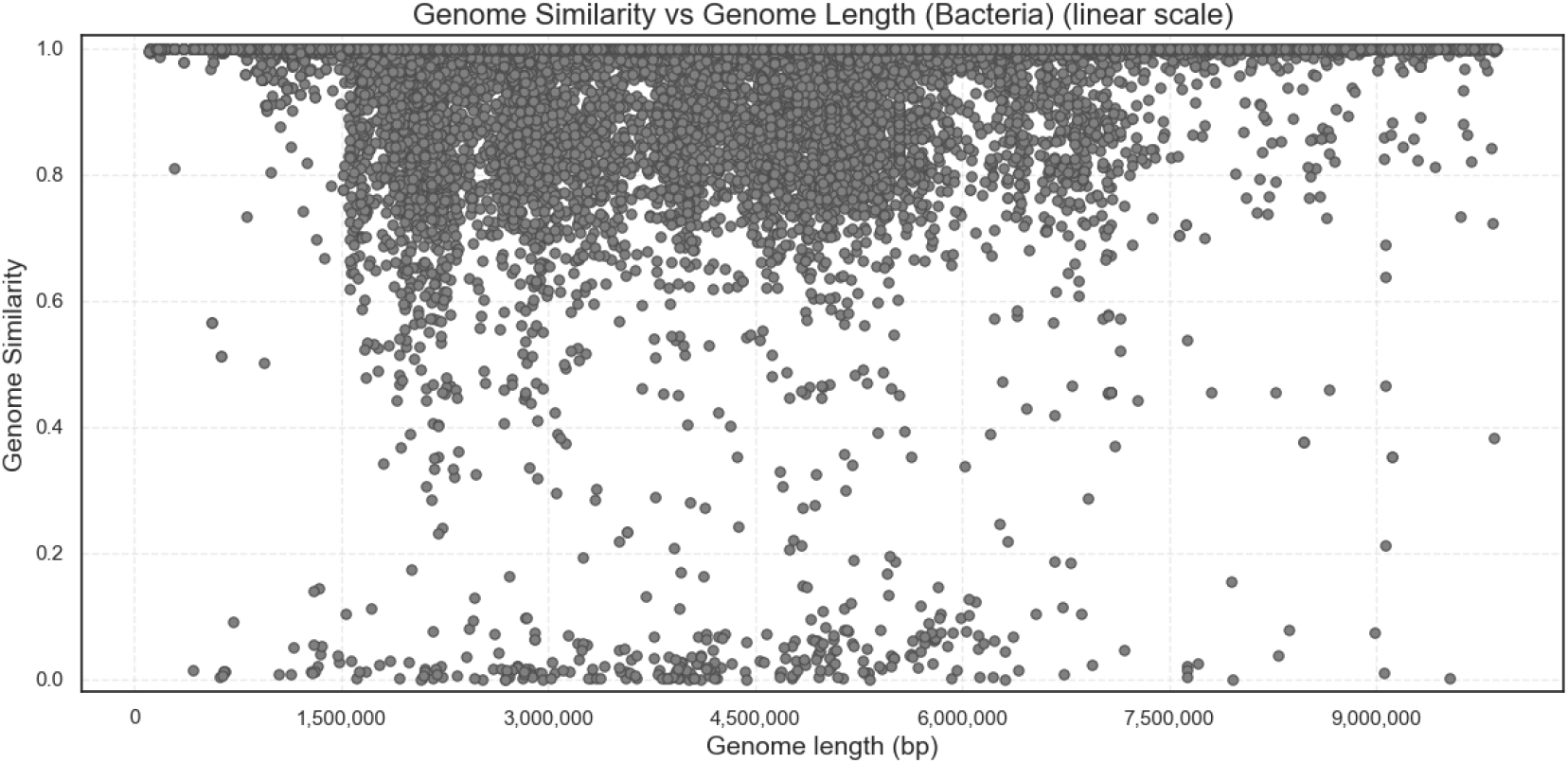
Relationship between genome similarity and genome length in bacterial assemblies. Each point represents a pairwise comparison between bacterial genome assemblies matched at the strain level across reference databases. The x axis shows genome length in base pairs on a linear scale, and the y axis shows the calculated genome similarity for each comparison. Most genome pairs cluster near complete similarity across a broad range of genome lengths, while a subset of comparisons forms a long tail of lower similarity values. No clear systematic relationship between genome length and similarity is apparent, indicating that genome size alone does not explain the observed variability in cross database concordance.

**Figure S9.**
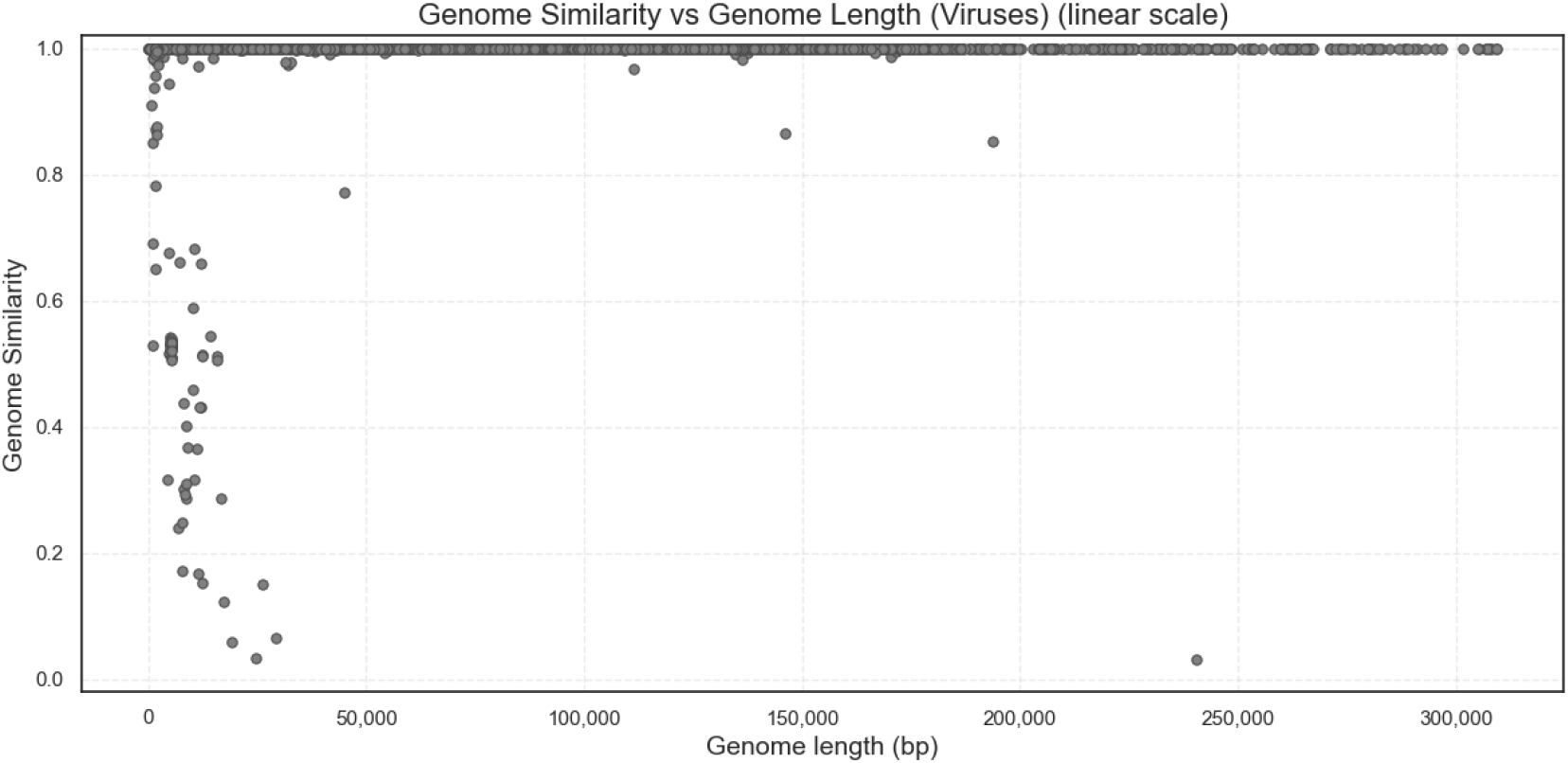
Relationship between genome similarity and genome length in viral assemblies. Each point represents a pairwise comparison between viral genome assemblies matched at the species level across reference databases. The x axis shows genome length in base pairs on a linear scale, and the y axis shows the calculated genome similarity for each comparison. Most viral genome pairs cluster near complete similarity across a wide range of genome lengths, reflecting strong cross database concordance. A limited number of comparisons form a tail of lower similarity values, but no clear systematic relationship between genome length and similarity is observed, indicating that genome size alone does not account for the variability in agreement across databases.

**Figure S10.**
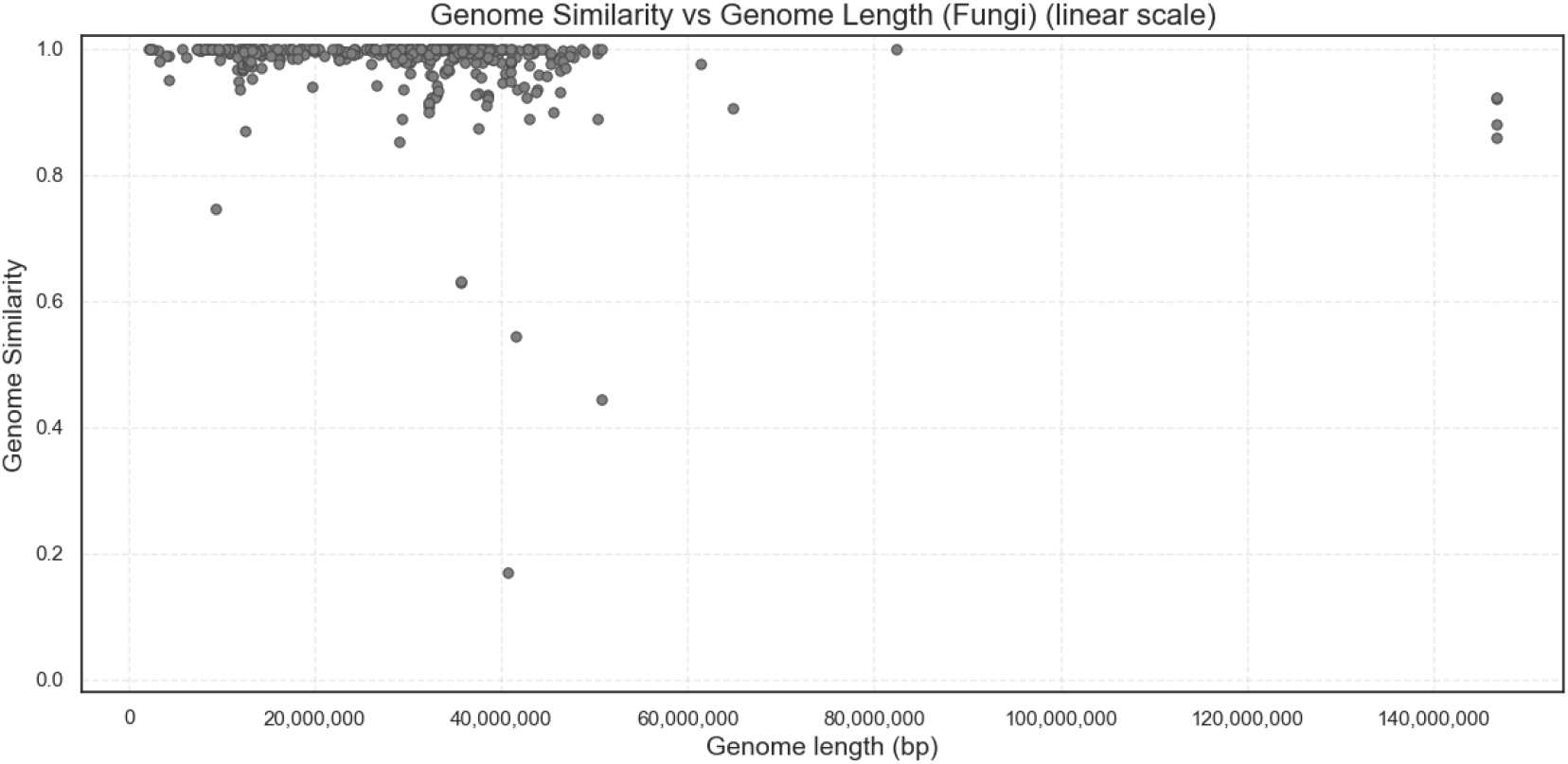
Relationship between genome similarity and genome length in fungal assemblies. Each point represents a pairwise comparison between fungal genome assemblies matched at the species level across reference databases. The x axis shows genome length in base pairs on a linear scale, and the y axis shows the calculated genome similarity for each comparison. Most genome pairs cluster near complete similarity across a broad range of genome lengths, with a limited number of lower similarity outliers. No clear systematic relationship between genome length and similarity is observed.

**Figure S11.**
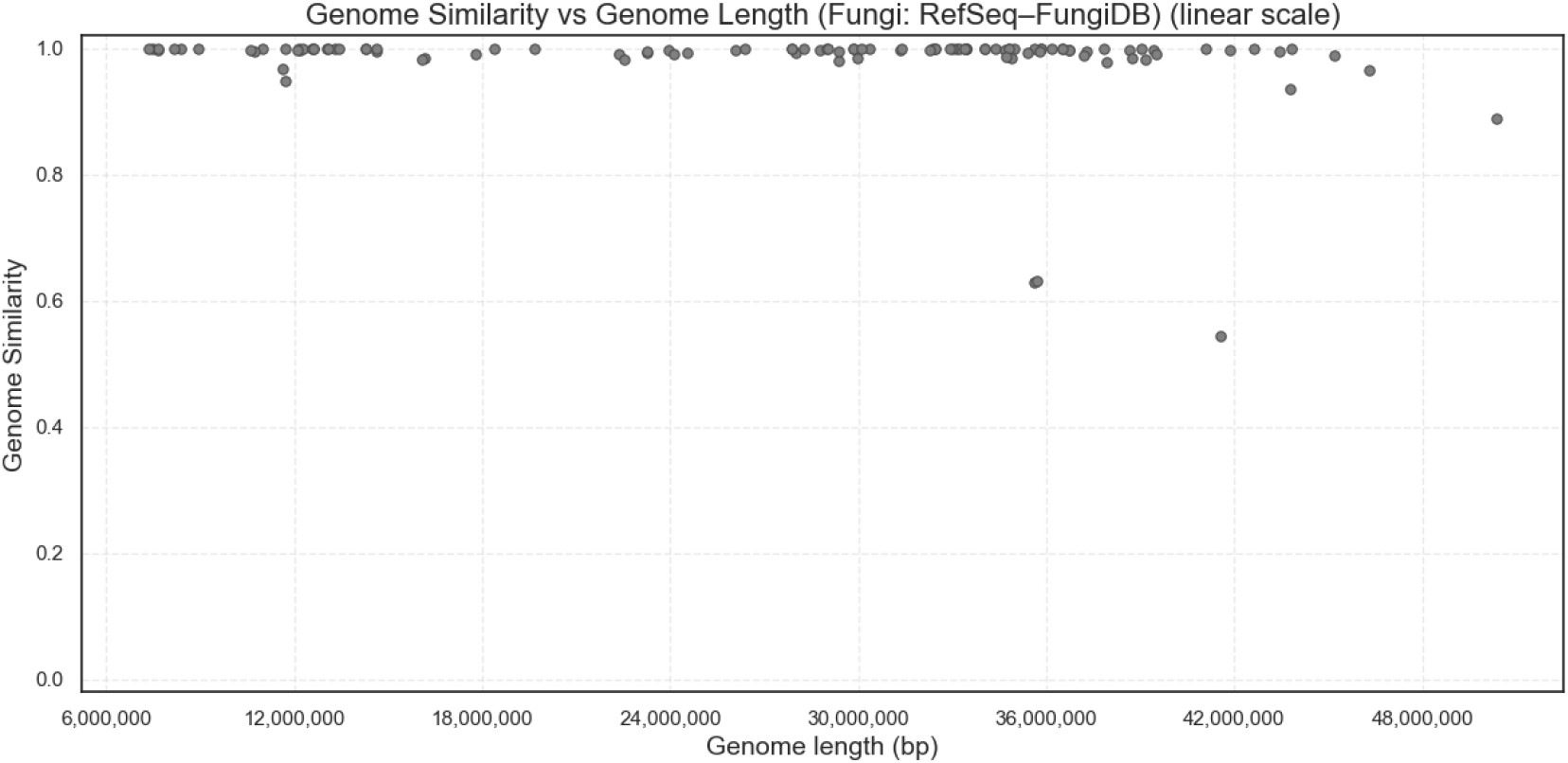
Relationship between genome similarity and genome length in fungal assemblies: RefSeq–FungiDB comparisons. Each point represents a species level pairwise comparison between fungal genome assemblies obtained from RefSeq and FungiDB. Genome similarity is plotted against genome length on a linear scale. The majority of comparisons show high similarity across the observed genome length range, with a small subset of lower similarity values. Variation does not appear to follow a consistent dependence on genome size.

**Figure S12.**
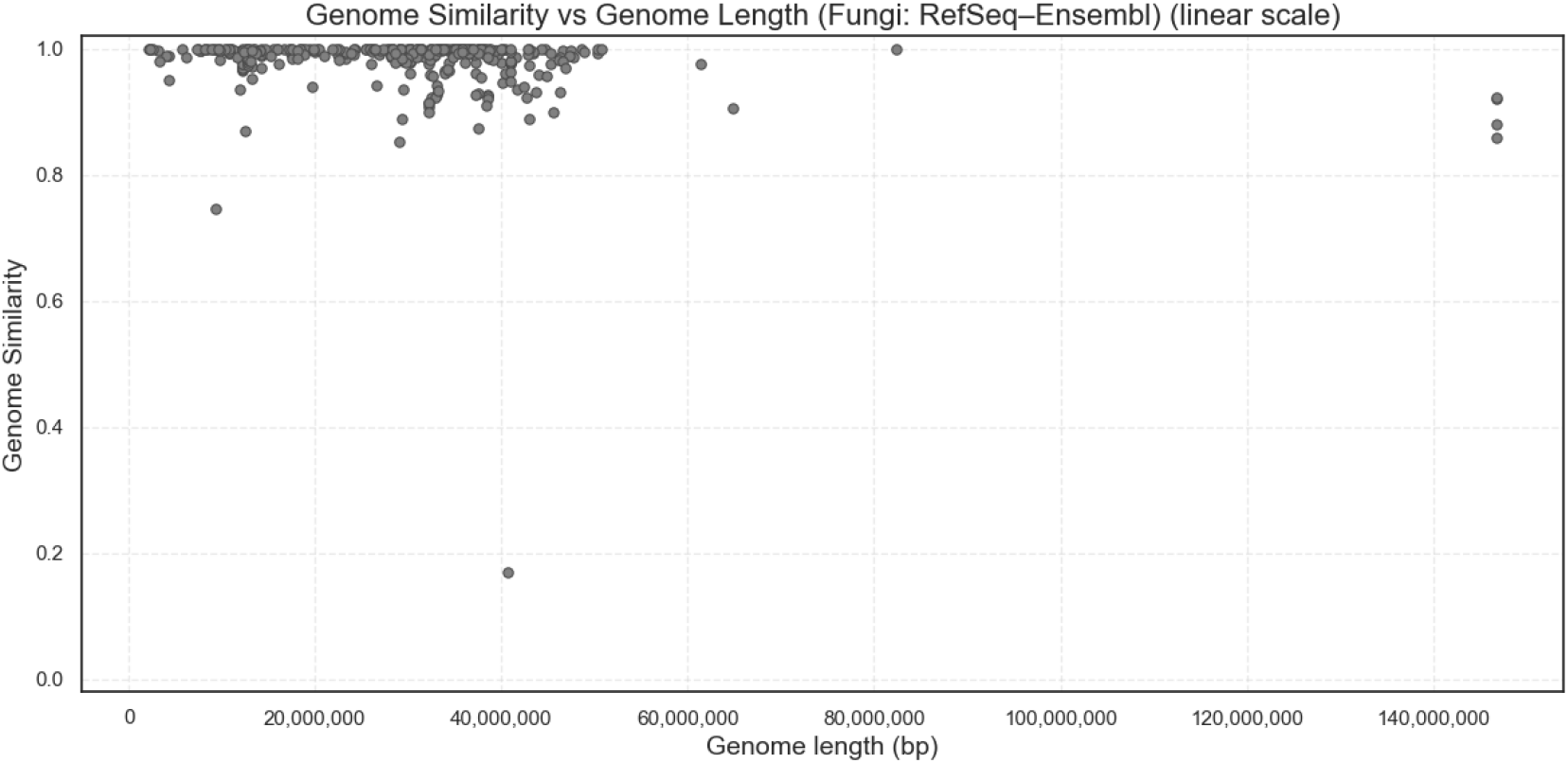
Relationship between genome similarity and genome length in fungal assemblies: RefSeq–Ensembl comparisons. Each point represents a species level pairwise comparison between fungal genome assemblies obtained from RefSeq and Ensembl. Genome similarity is plotted against genome length on a linear scale. Most comparisons cluster near complete similarity across diverse genome lengths, with a limited number of lower similarity outliers. As in the other fungal analyses, genome length does not exhibit a clear systematic association with similarity.

**Supplementary Table S1.** Tool comparison on a known ground truth.

| Tool | Seq length<br>s (bp) | Total length<br>h (bp) | Alignment<br>length<br>(bp) | Matches* | Mismatches | Indels | Unaligned (bp) |
| --- | --- | --- | --- | --- | --- | --- | --- |
| Ground truth | 1700<br>(Ref)<br>/<br>1701<br>(Qry) | 3401 | 1203<br>(Ref) /<br>1203<br>(Qry) | 1177 | 21 | 5 | 497<br>(Ref) /<br>498<br>(Qry) |
| BLAST | 1700<br>(Ref)<br>/<br>1701<br>(Qry) | 3401 | 1203<br>(Ref) /<br>1203<br>(Qry) | 1177 | 21 | 5 | 497<br>(Ref) /<br>498<br>(Qry) |
| MUMmer4 | 1700<br>(Ref)<br>/<br>1701<br>(Qry) | 3401 | 1194<br>(Ref) /<br>1197<br>(Qry) | 1170 | 21 | 2 | 506<br>(Ref) /<br>504<br>(Qry) |
| GS-Align | 1700<br>(Ref)<br>/<br>1701<br>(Qry) | 3401 | 1194<br>(Ref) /<br>1194<br>(Qry) | 980 | 233 | 32 | 506<br>(Ref) /<br>507<br>(Qry) |

**Supplementary Table S2.** Distribution of bacterial genomes by number of contigs shared between RefSeq and BV-BRC. Each row represents the number of genomes (cases) that have the same number of contigs (from 1 to 10) in both databases.

| Number of contigs in both databases | Number of cases |
| --- | --- |
| 1 | 21110 |
| 2 | 9084 |
| 3 | 5815 |
| 4 | 4073 |
| 5 | 3109 |
| 6 | 2334 |
| 7 | 1763 |
| 8 | 1424 |
| 9 | 1389 |
| 10 | 1345 |

